# The dynamics of spawning acts by a semelparous fish and its associated energetic costs

**DOI:** 10.1101/436295

**Authors:** Cédric Tentelier, Colin Bouchard, Anaïs Bernardin, Amandine Tauzin, Jean-Christophe Aymes, Frédéric Lange, Charlotte Recapet, Jacques Rives

## Abstract

1. During the reproductive season, animals have to manage both their energetic budget and gamete stock. In particular, for semelparous capital breeders with determinate fecundity and no parental care other than gametic investment, the depletion of energetic stock must match the depletion of gametic stock, so that individuals get exhausted just after their last egg is laid and fertilized. Although these budgets are managed continuously, monitoring the dynamics of mating acts and energy expenditure at a fine temporal scale in the wild is challenging.
2. This study aimed to quantify the individual dynamics of spawning acts and the concomitant energy expenditure of female Allis shad (*Alosa alosa*) throughout their mating season.
3. Using eight individual-borne accelerometers for one month, we collected tri-axial acceleration, temperature, and pressure data that we analysed to i) detect the timing of spawning acts, ii) estimate energy expenditure from tail beat frequency and water temperature, and iii) monitor changes in body roundness from the position of the dorsally-mounted tag relative to the vertical plane.
4. Female shad had a higher probability to spawn during warmer nights, and their spawning acts were synchronized (both individually and inter-individually) within each active night. They experienced warmer temperature, remained deeper, swan more slowly and spent less energy during daytime than night time. Over one month of spawning, they performed on average 15.75 spawning acts, spent on average 6 277 kJ and died with a significant portion of residual oocytes. The acceleration-based indicator of body roundness was correlated to condition coefficient measured at capture, and globally decreased through the spawning season, although the indicator was noisy and was not correlated to changes in estimated energy expenditure.
5. Despite significant individual variability, our results indicate that female shad exhausted their energetic stock faster than their egg stock. Water warming will increase the rate of energy expenditure, which might increase the risk that shad die with a large stock of unspent eggs. Although perfectible, the three complementary analyses of acceleration data are promising for *in situ* monitoring of energy expenditure related to specific behaviour.

## Introduction

The energy acquired by living organisms is allocated to survival and reproduction, and natural selection is expected to favour optimal allocation, resulting in life-histories that maximize Darwinian fitness in the environment where evolution occurs (Pianka 1976; Stearns 1992; Roff 1993). When adult survival is low compared to juvenile survival, extreme reproductive effort (in gametogenesis, mating behaviour and parental care) can be selected and semelparity may arise, in which individuals die after their first and only breeding season. Semelparity is often accompanied with capital breeding, so that reproduction relies on energy reserves constituted before the breeding season (Bonnet et al. 1998). Although semelparity is sometimes quoted as “big bang reproduction”, its broad definition encompasses cases where individuals breed in several bouts within a breeding season (Kirkendall and Stenseth 1985; Hughes 2017). Furthermore, in species with no other parental care than gametic investment, the optimal allocation should result in individuals dying just after their last progeny has been produced, and deviation from optimality consists in individuals either surviving after their last egg has been laid and fertilized, or dying of exhaustion while still bearing unlaid eggs (Heimpel and Rosenheim 1998). Hence, the schedule of breeding events and the dynamics of energy expenditure during the breeding season, which may be pronounced in semelparous capital breeders, is a crucial aspect of reproductive strategy.

Beside energy expenditure, the temporal distribution of breeding events within a season may be linked to ecological, pheromonal or behavioural cues that synchronize breeding activity (Gochfeld 1980; Ims 1990; Fürtbauer et al. 2011). Here, internal and external stimulations probably interact, with the hormone-driven reproductive cycle and energetic condition determining whether an individual is ready to breed at a given time, and social or ecological factors triggering the actual breeding behaviour (Jovani and Grimm 2008; Koizumi and Shimatani 2016). At the ultimate level, reproductive synchrony affects intraspecific competition for resources (including mates; Emlen and Oring 1977) and the risk of predation on breeders or their offspring (Ims 1990).

Tracking the schedule of breeding events and the dynamics of energy expenditure of breeding individuals in the wild is technically challenging. The timing of breeding events along the breeding season requires thorough observation of repeatedly detectable individuals, for example through video recording of highly sedentary individuals (e.g. Borgia 1985). Energy expenditure has been monitored at the population level, by quantifying energy reserves on different individuals sampled at different stages of the breeding season (e.g.

Hendry and Berg 1999). In some cases, the same individuals were captured at the beginning and at the end of the breeding season, and the energy expenditure quantified with difference in mass (Anderson and Fedak 1985; Rands et al. 2006), body composition (Hendry and Beall 2004; Casas et al. 2005), or concentration of plasma metabolites (Gauthey et al. 2015). Likewise, in terrestrial animals, the turnover of 18O and ^2^H from doubly labelled water quantifies the average field metabolic rate at the individual level between two sampling occasions (Nagy et al. 1999). All these methods give a good idea of the total energy expenditure over the whole breeding season but often lack the temporal resolution to document the longitudinal dynamics of energy expenditure all along the breeding season.

With the advent of bio-logging, the temporal resolution of individual data collected in the field has tremendously increased. Among bio-loggers, accelerometers, often coupled with other sensors such as thermometers, have been increasingly used, and advances in their energetic efficiency now allows to record high-frequency data for weeks or months, making them valuable tools for longitudinal studies of breeding behaviour in many organisms. In this context, acceleration data can quantify both the overall movement of tagged individuals (overall dynamic body acceleration), which can be translated in energy through laboratory calibration (Wilson et al. 2006; Groscolas et al. 2010; Collins et al. 2016; Hicks et al. 2017; Wilson et al. 2019), and the number and temporal distribution of key behaviours such as mating events, if these produce a typical acceleration pattern (Tsuda et al. 2006; Brown et al. 2013; Sakaji et al. 2018).

This study aimed at describing the activity and change in body condition of female Allis shad (Alosa alosa L. 1758) throughout their spawning season. Allis shad is a semelparous anadromous clupeid, a capital breeder with determinate fecundity (Baglinière et al. 2003). It is also a major halieutic resource in the Atlantic coast of southern Europe, and the size of most of its populations has plummeted since the beginning of the 21st century. Its economical and conservation value and its spawning behaviour are such that the timing of its reproduction is very well documented at the population level. The spawning act consists of at least one male and one female swimming side by side during five to ten seconds while describing three to five circles of one meter in diameter and beating the water surface vigorously with their tail (Baglinière and Elie 2000). The typical splashing noise (35 dB at one meter) can be heard and recorded from the river bank, and the number of splashes recorded during the spawning season is often used as an indicator of the number of spawners in a population (Chanseau et al. 2004). Despite the wealth of data on the timing of spawning acts across the season and within nights at the population level, no reliable data exists at the individual level (but see Acolas et al. 2004 for a previous attempt), which is the appropriate level to eventually address the fitness associated to behavioural strategies.

Using accelerometers, we investigated three main aspects and tested specific predictions:

1. We described the individual schedule of spawning events, which were identified using a characteristic acceleration pattern. We tested whether the probability of spawning increased solely with water temperature (Paumier et al. 2019) or was also affected by internal and social factors. In particular, as oocytes seem to mature in five to seven batches (Cassou-Leins and Cassou-Leins 1981), we expected shad to perform five to seven spawning acts separated by a consistent time lapse corresponding to the maturation time of each batch. Moreover, given the aggregative behaviour of shad, which shoal even during the spawning season (Baglinière and Elie 2000), we tested whether different females synchronized their spawning acts within a night.
2. We quantified global activity as tail beat frequency, and converted it to energy expenditure through an energetic model including water temperature. This was used to identify variability in energy expenditure between periods of the spawning season and to test the effect of energy management on the risk of dying with unlaid eggs. In particular, we tested the predictions that shad were more active and spent more energy during the night than during the day, and that the individuals that better managed their energy expenditure (higher night time energy expenditure and lower daytime energy expenditure) died with fewer residual eggs.
3. We tested the possibility to use the angle between the dorsally-mounted accelerometer and the vertical plane as an indicator of body roundness of the fish. This novel method would allow monitoring the slimming process through the spawning season, and relating it to periods of high energy expenditure. We predicted that if the angle of the accelerometer was a reliable indicator of fish condition, it should a) be correlated to the individual coefficient of condition measured at the beginning and the end of the spawning season, b) decrease across the spawning season, as the fish slims, c) decrease proportionally to mass loss, d) decrease more rapidly during periods of high energy expenditure.

## Methods

### Characteristics of the species studied

Allis shad (*Alosa alosa* L.) is an anadromous clupeid fish distributed along the Atlantic coast of Europe, from Portugal to the British Isles, with the main populations dwelling in the French rivers Garonne, Dordogne and Loire (Baglinière and Elie 2000). Across its distribution, Allis shad is considered as a semelparous species, although spawning marks on scales suggest that a very small proportion of individuals may spawn on two consecutive years (Mennesson-Boisneau and Boisneau 1990; Taverny 1991). After having spent a few months in freshwater as juveniles and four to six years at sea, shad undertake freshwater upstream migration during which they fast, thus qualifying as capital breeders. Gonad maturation occurring during migration leads females to bear between 13 000 and 576 000 eggs, reaching an average gonad mass of 221 g and a gonad / somatic mass ratio of 15% on spawning grounds (Cassou-Leins and Cassou-Leins 1981; Taverny 1991). Spawning typically occurs at night in a 0.3 to 3-metre-deep glide. As mentioned above, the spawning act consists of rapid circular swimming and vigorous tail beating for five to ten seconds, and should produce an easily detectable acceleration pattern. Although very few data are available, the number of splashes recorded upstream from dams where migrating shad were counted suggest that females perform on average five to twelve spawning acts during the season (Fatin and Dartiguelongue 1996; Acolas et al. 2006). Moreover, based on the dynamics of ovary index and oocyte diameter measured on individual caught and dissected across the season, Cassou-Leins & Cassou-Leins (1981) postulated that shad mature their eggs in five to seven batches.

The spawning season spans from early-May to late July, but dead individuals can be collected downstream from spawning ground as soon as late May. The number of splashes heard on spawning grounds increases with increasing water temperature (Baglinière and Elie 2000; Paumier et al. 2019), but this could be due to either more individuals being sexually active or individuals to be more active at higher temperature. Within a night, the temporal distribution of splashes in large populations typically follows a Gaussian distribution centred on 2 AM and spanning from 10 PM to 6 AM (Cassou-Leins and Cassou-Leins 1981).

Shad being capital breeders, samples of different individuals caught at different times during the migration and reproduction have shown dramatic changes in somatic mass and tissue composition (Cassou-Leins and Cassou-Leins 1981; Bengen 1992), but no longitudinal data exist at the individual level.

### Fieldwork

We conducted this study in spring 2017 and 2018 in the Nivelle, a 39 km long coastal river situated in the Northern Basque Country, France, and draining a 238 km^2^ basin. The downstream limit of the study zone was the impassable Uxondoa weir, situated 12 km upstream from the river mouth (43°21’40.64”N, 1°35’13.99”W), and equipped with a vertical slot fishway and a trap where a yearly average of 230 (min = 26; max = 688) migrating shad have been counted since 1996, with less than 30 individuals per year since 2015. Five kilometres upstream from Uxondoa stands another impassable weir equipped with a fishway and a trap, where shad have almost never been captured.

The Uxondoa fish trap was controlled daily throughout spring (ECP 2018). Nine female shad were captured and tagged in 2017, and 15 in 2018. All experimental procedures comply with French and European legislation, and were approved by the legal representative (prefectural decree #64-2017-04-25-004) and the ethical committee for birds and fishes in the French region Nouvelle Aquitaine (authorization #2016020116037869). The tagging procedure was quite similar to Breine (2017) for twaite shad (*Alosa fallax*). We anesthetized each individual in a bath of 15 mg/L benzocaine diluted in river water before weighing it, measure its fork length, gently press the abdomen to check the absence of sperm emission, and finally tag it with a radio transmitter emitting at a unique frequency (F2020, ATS, Isanti, MN, USA) and a three-dimensional accelerometer (WACU, Atesys-Montoux, Huguenau, France). For tagging, each fish was kept in a 20 L tank filled with anaesthetic solution (Fig. 1.A). A water pump placed in the fish’s mouth and a stone bubbler in the bath ensured a good circulation of aerated water during the tagging procedure. Adjustable plastic plates covered with foam were placed vertically against both flanks to maintain the fish in an upright position, with only its back being above water surface. The radio transmitter and the accelerometer were cleaned with a povidone iodine solution (Betadine®) and dried with surgical cotton. The two Teflon-coated metallic wires of the radio transmitter were inserted 1 cm under the dorsal fin through sterile hollow needles. The hollow needles were then removed, and the metallic wires were passed through holes drilled in the accelerometer’s lug, secured with plastic eyelets and aluminium sleeves, and the extra length was cut. The radio transmitter and the accelerometer weighed 8.6 g and 9 g, respectively, so the weight was balanced on both sides of the fish. After tagging, each fish was placed in a 50 L box filled with river water, and released upstream from the weir upon waking.

**Figure 1.**
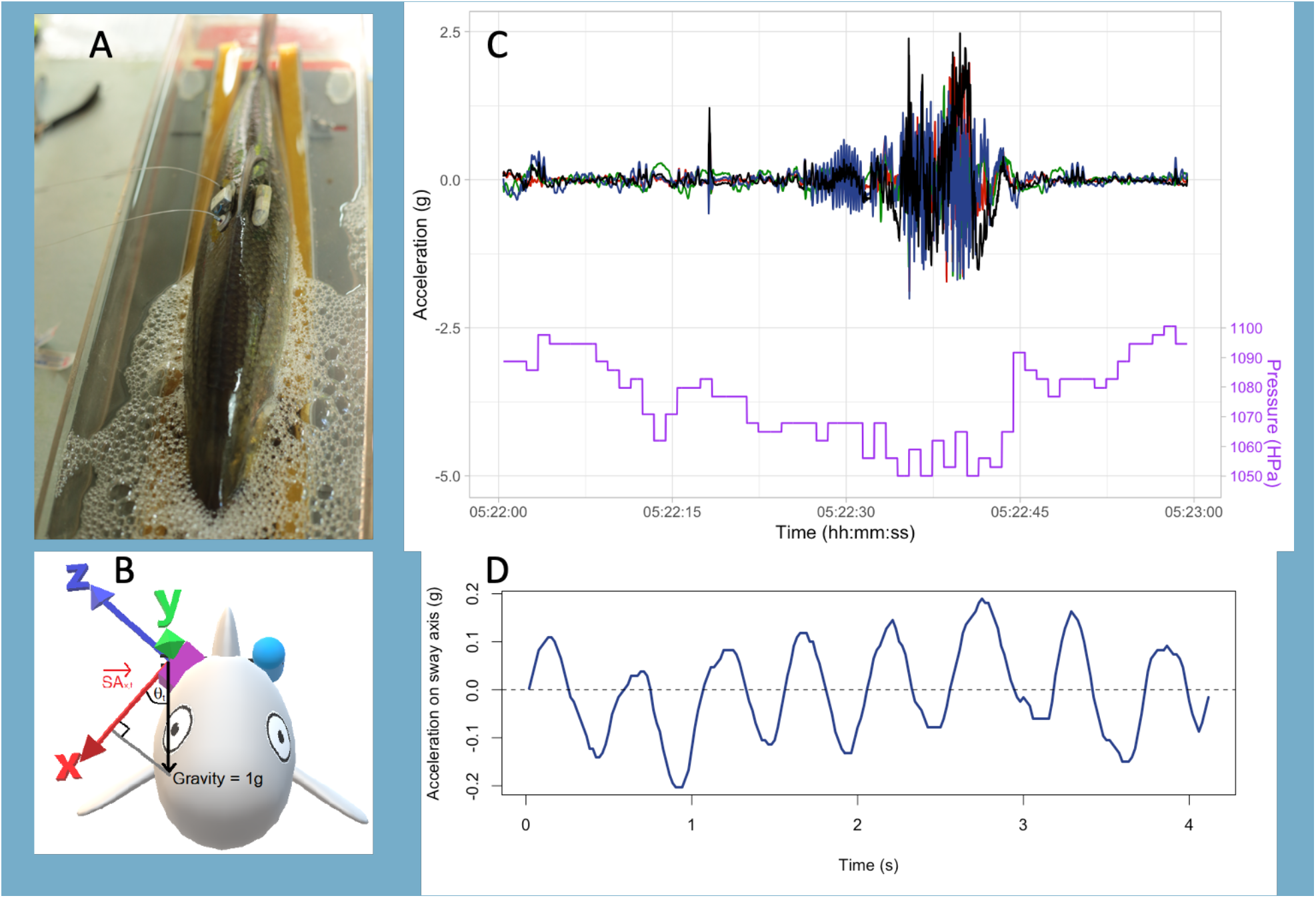
**A.** Allis shad were tagged under anaesthesia with a radio transmitter on the left flank and an accelerometer on the right flank. The accelerometer continuously recorded acceleration on three axes. **B.** The static component of acceleration recorded on the x-axis at time *t*, 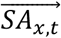, corresponds to the orthogonal projection of gravity, which is vertical by definition, on this axis. Hence, the angle *θ* between the x-axis and the vertical plane can be computed for any time *t* as 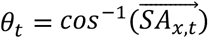. **C.** The dynamic component of acceleration (here smoothed with a 31-point wide median filter) can be used to detect spawning acts, characterized by increased acceleration on both x (red), y (green) and y (blue) axes, resulting in a peak in the norm of the 3-dimensional acceleration vector (black), and accompanied by a decrease in hydrostatic pressure (purple). **D.** After correction for the *θ* angle, the z-axis corresponds to the sway axis of the fish, and its dynamic component (here smoothed with a 5-point wide median filter) can be used to compute tail beat frequency (TBF) as the number of zero-crossings per unit of time.

From its release at Uxondoa weir until its death at the end of the spawning season, each tagged shad was localized twice a day using a mobile receiver (R2100, ATS, Isanti, MN, USA) and a loop antenna, in order to check whether it was still alive and to obtain its position. The radio transmitters were set to double their pulsing rate after 8 h of total immobility. Within two days after double pulse was detected, the dead fish and the tags were recovered by snorkelling, and the whole fish and its ovaries were immediately weighed.

### Processing acceleration data

The loggers used in this study recorded acceleration between −8 g and +8 g (1 g = 9.81 m.s^−2^) in each of the three dimensions at an average frequency of 50 logs per second. Every second, they also recorded the temperature, pressure, date, time and exact number of acceleration logs within the past second. Both dynamic and static (gravitational) acceleration were used to estimate three types of variables: body roundness as the angle of the accelerometer with the vertical plane, tail beat frequency and occurrence of spawning acts.

The static, gravitational, component on each axis *i* (x, y and z) was extracted from the raw signal at each time *t* by replacing each data point 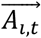 by the average of the points within a window of width *w* centred on *t*, so that:

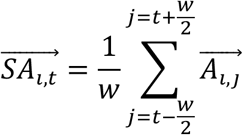

Unlike what is usually done (Brown et al. 2013), these data were not used to assess the posture of the fish, as no known behaviour in shad involves change of posture. Instead, we hypothesized that it may reflect the shape of the fish. Indeed, since the accelerometer was pinned on the dorsal part of the fish’s flank, the angle *θ* between the x-axis of the tag and the vertical plane may be linked to the roundness of the fish, a proxy of its energetic reserves. This angle could be computed at any data point of static acceleration, as 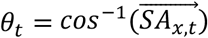 where *θ_t_* is the angle at time *t* and 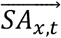 is the static acceleration computed on the x-axis at time *t* (Fig. 1.B). To track change in body roundness through the season, *θ* was estimated for each fish from 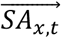 computed over *w*=180 000 points (i.e. 1 h at a 50Hz sampling frequency) every eight hours (6AM, 2PM, 10PM), from its release until its death.

The dynamic component of acceleration was computed at every time point on each dimension 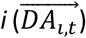 as the raw signal minus the static acceleration computed over *w*=250 points (= 5 s at 50 Hz). This was used to count the number of spawning events performed by each tagged shad. To do this, the acceleration pattern typical of spawning act was first described in a controlled experiment and then searched in the field-collected data. To characterize the typical pattern associated with spawning act, a male (420 mm, 855 g) and a female (460 mm, 1140 g) shad were captured at Uxondoa trap, tagged as described above, and observed for one month (June 2016) in a basin (400 m^2^, 0.6 m water depth) at the INRAE experimental facilities in Saint Pée sur Nivelle (ECP 2018). Eight spawning acts were recorded on video or audio, which could be compared with acceleration data. The typical pattern of acceleration associated to the spawning act was a rise in the norm of the 3D acceleration vector, which stayed above 3 g for at least 3 seconds, mainly driven by acceleration on the z-axis corresponding to TBF reaching up to 15 beats per second. (Fig. 1.C). A decrease of hydrostatic pressure was typically associated to this pattern, due to the fish reaching the water surface during the spawning act. These criteria for identification of spawning acts were robust, as they were always detected when spawning was visually observed (no false negative) and never observed when spawning was not visually observed (no false positive). These criteria were implemented in an algorithm to scan the accelerograms collected in the field in 2017. This automated identification was validated by visual comparison of accelerogram sequences automatically identified in 2017 with sequences of eight acknowledged spawning acts recorded in 2016.

The dynamic acceleration was also used to quantify the activity of the fish through Tail Beat Frequency (TBF), computed as the number of zero-crossings per second of 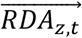 on which a median filter (window width = 5 points) was applied for noise reduction (Fig. 1.D). Acceleration on the sway axis is commonly used to measure TBF on fishes of different species and sizes (Kawabe et al. 2003; Tsuda et al. 2006; Broell and Taggart 2015; Brownscombe et al. 2018), and preliminary data on the two captive individuals mentioned above showed that our acceleration-based measurement of TBF corresponded to the visual measurement. TBF can be computed at every instant, but for further analysis we used average computed for every minute.

We estimated the energy consumption for every minute of each individual’s reproductive season, using its tail beat frequency (computed as described above), the temperature recorded by the individual logger, and equations derived from the data of Leonard et al. (1999) and Castro-Santos & Letcher (2010) on American shad, *Alosasa pidissima*. Leonard et al. (1999) measured oxygen consumption and tail beat frequency of 18 American shad in a swim tunnel respirometer with water temperature varying between 13°C and 24°C. On their data, we fitted a linear mixed model (lmer function in lme4 package; Bates et al. 2014) with individual random intercept and fixed effects of temperature and TBF, on log-transformed oxygen demand (MO_2_, in mmol O_2_.kg^−1^.h^−1^). Then, assuming that 0.4352 kJ of somatic energy are burnt per mmol O_2_ (Brett and Groves 1979), we used the three parameters of the mixed model (average intercept = 1.0064, slope of temperature = 0.0531, and slope of TBF = 0.2380) to compute the amount of energy *E_i,t_* (in kJ) consumed by each fish *i* for each minute *t* of its spawning period, from its body mass *M_i,t_* (in kg), the average temperature recorded by its logger during that minute *T_i,t_* (in °C), and its average tail beat frequency during that minute *TBF_i,t_* (in beat per second):

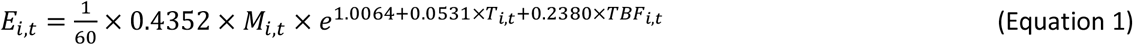

Since energy consumption continually reduces fish mass, we iteratively modelled fish mass and energy expenditure at each minute of the spawning season. For this, we assumed that shad get 69% of their energy from lipids and 31% from proteins to fuel their metabolism (Leonard & McCormick, 1999), and that one gram of fat yields 39.54 kJ, against 23.64 kJ per gram of protein (Craig et al. 1978), so fish mass at the *t*^th^ minute was:

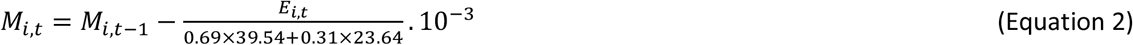

Finally, the exact timing of each individual’s death was detected as a clear change in the acceleration signal, where the gravitational component indicated that the fish switched from upright posture to lying on its side or upside down, and the only dynamic component left was the tenuous jiggling due to water flowing over the tag.

Beside acceleration, the pressure data given by the tags was processed to translate it to depth by computing hydrostatic pressure (1hPa.cm^−1^) as the difference of pressure recorded every second by each tag and the atmospheric pressure recorded every 20 minutes by a meteorological station situated at the INRAE facilities, less than one kilometre from the spawning ground.

### Statistical analysis

All analyses were performed on R (R Development Core Team 2008). A zero-inflated mixed regression (package glmmTMB; Brooks et al. 2017) was used to test the effect of temperature on the number of spawning acts performed by each individual on each night. The Binomial component of the model informed on the effect of temperature on the probability to perform any spawning act, while the Poisson component tested the effect of temperature on the number of spawning acts. Individual was used as a random effect on both components. To test whether females spawning in the same night synchronized their spawning acts with each other, we performed 10 000 permutations of the hour of spawning acts recorded along all nights, and compared the median of the delay to the nearest spawning act for actual data and for simulated data. Because our aim was not to test the synchrony between acts performed by the same female, only the first act performed by each female on a given night was considered in this analysis.

As mentioned above, TBF was computed for every minute of each individual’s spawning season, to track its energy expenditure. As an indicator of fish activity, average TBF was also aggregated over eight-hour time windows corresponding to three periods of the day: morning (6AM-2PM), afternoon (2PM-10PM) and night (10PM-6AM). The limits of the night period correspond to the earliest and latest hour of spawning activity classically recorded on spawning grounds (Cassou-Leins and Cassou-Leins 1981). To test whether shad adopted a different behaviour at different periods of the day, linear mixed models with individual random intercept and period of the day as a fixed effect were fitted to TBF, temperature and depth. For TBF, we also added a binary covariate indicating whether the fish had been tagged for more or less than three days, to test whether recent tagging increased activity (as observed in Atlantic salmon by Føre et al. 2020). All mixed models were fitted using the package lme4 for R (Bates et al. 2014), significance of fixed effects was tested using the likelihood ratio test (LRT χ^2^) between the model including the fixed effect and the nested model excluding it. Marginal and conditional R^2^ (R^2^_m_ and R^2^_c_), indicating the proportion of variance explained by the fixed effects and by the whole model, respectively, where computed using R package MuMIn (Barton 2009).

To our knowledge, static acceleration has never been used to assess changes in animal body roundness. As mentioned in the introduction, we tested four predictions resulting from the hypothesis that the angle *θ* between the x-axis of an individual’s accelerometer and the vertical plane was an indicator of body roundness. First, we fitted a Pearson correlation between Fulton condition coefficient (100*mass (in grams)/length (in centimetres)^3^) and *θ,* both measured at the initial capture and at death. Second, a linear mixed model, with individual random intercept and slope was used to test whether *θ* decreased with time since individual release. Depth was added in this model, to account for the possibly positive effect of swim bladder inflation (hence negative effect of depth) on body roundness. Third, to test if change in *θ* over the breeding season was proportional to change in condition, a correlation was performed between the differences in *θ* and in Fulton condition coefficient from release to death. Finally, to test whether the decrease in *θ* could be linked to temperature, activity, energy expenditure or the moment of the days, we fitted linear mixed models with the difference in *θ* between the first and the last hour of each 8-hour period and an individual random intercept. Four models were tested, 1) the null model with only random intercept, 2) a model with average temperature and TBF during the 8-hour period as fixed effects, 3) a model with cumulated energy expenditure during the 8-hour period as a fixed effect, and 4) a model with the moment of the period (morning, afternoon, night) as a fixed effect. To compare mixed models corresponding to alternative hypotheses, we used Akaike Information Criterion corrected for small sample bias (AICc) on R package MuMIn (Barton 2009).

The R code for statistical analysis and the data sets on which they were performed are available in the institutional data repository of the INRAE (French National Institute for Agriculture Food and Environment): https://doi.org/10.15454/NTFYCC.

## Results

The nine female shad tagged in 2017 survived between 20 and 37 days (mean=26 days), and all tags were retrieved within one or two days after fish died, although one tag stopped recording data after ten days. The 2018 campaign was much less successful: two fish died one week after tagging, before any spawning acts were recorded; two fish lost their tags three and four days after tagging. The eleven remaining fish were radio tracked throughout the spawning season, until two exceptional floods (on June 7^th^ and 16^th^) flushed them down to the estuary and the ocean where high water conductivity prevented further radio tracking, hence making tag retrieval impossible. Eventually, among the 25 tagged shad, only eight fully exploitable and one partially exploitable accelerograms could be collected. This sample of eight females represented half of the females that passed the Uxondoa weir in 2017, but the power of Spearman correlations to detect even a strong correlation of 0.5 between variables observed at the individual level was only 0.22. The minimal, median and maximal fork length and body mass of the eight females were 470, 510 and 550 mm, and 1320, 1635 and 1810 g.

According to their accelerograms, the eight female shad performed 7, 9, 12, 14, 14, 17, 24 and 26 spawning acts (mean=15.75). The total number of spawning acts performed by each female was correlated to none of the individual variables tested (Fig. 2).

**Figure 2.**
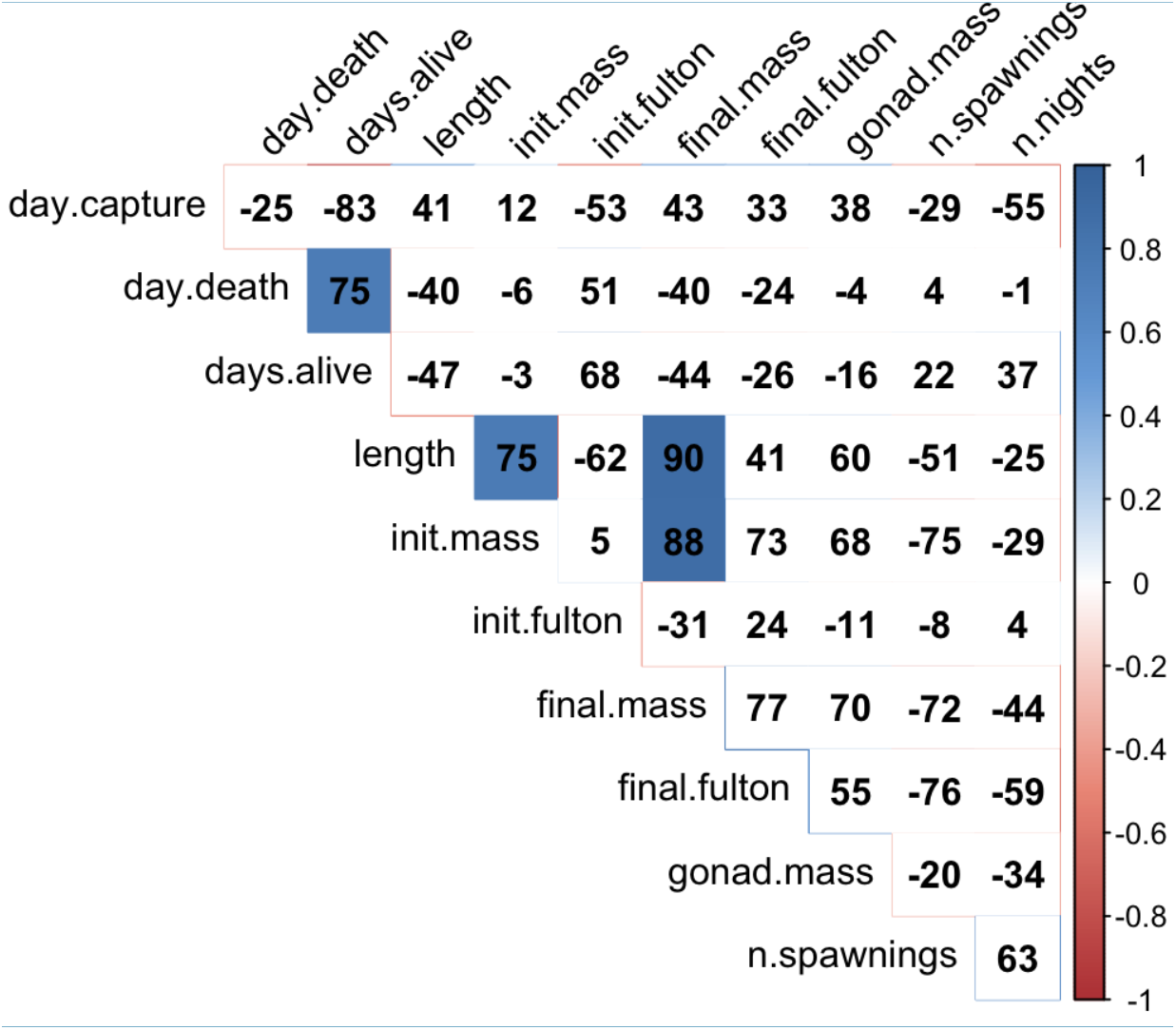
Spearman coefficient of correlation (expressed as percentage) between biometric and reproductive variables of female Allis shad. In order, day of capture, day of death, number of days between capture and death, body length, initial mass, initial Fulton coefficient of condition, final mass, final Fulton coefficient of condition, final gonad mass, number of spawning acts, number of nights with at least one spawning act.

For each female, spawning acts were distributed in three to six nights (mean=4.87) each separated by zero to eight nights (mean=3.55) without spawning acts, resulting in individual spawning seasons ranging from 14 to 24 nights (mean=18.12) from the first spawning act to the last (Fig. 3.A). The tagged females performed their first spawning act zero to eight days after tagging (mean=3). Only on eight occasions did a female perform a single spawning act during a night, so the 137 remaining acts were performed in volleys. At the individual level, an active night comprised from two to eight acts (mean=3.23) performed in two to 84 minutes (mean=29.7). The mixed zero-inflated Poisson regression indicated that water temperature during the night had a positive effect on the probability that a female performed at least one spawning act (negative effect on the zero inflation; z=−1.95, p=0.05; probability of a spawning act estimated 0.18 at 18°C and 0.23 at 19°C) but no effect on the number of spawning acts in the volley (the Poisson component; z=−1.44, p=0.15). Cumulated over all the season, the temporal distribution of spawning acts within the night followed a Normal distribution, centred on 3AM, with 95% of spawning acts occurring between 0:30AM and 5:30AM. However, the permutation test on the hour of spawning acts indicated synchrony between acts performed by different females on the same night: the median time lag to the nearest act was one hour for observed data, which corresponds to the first percentile of simulated data (Fig. 3.B).

**Figure 3.**
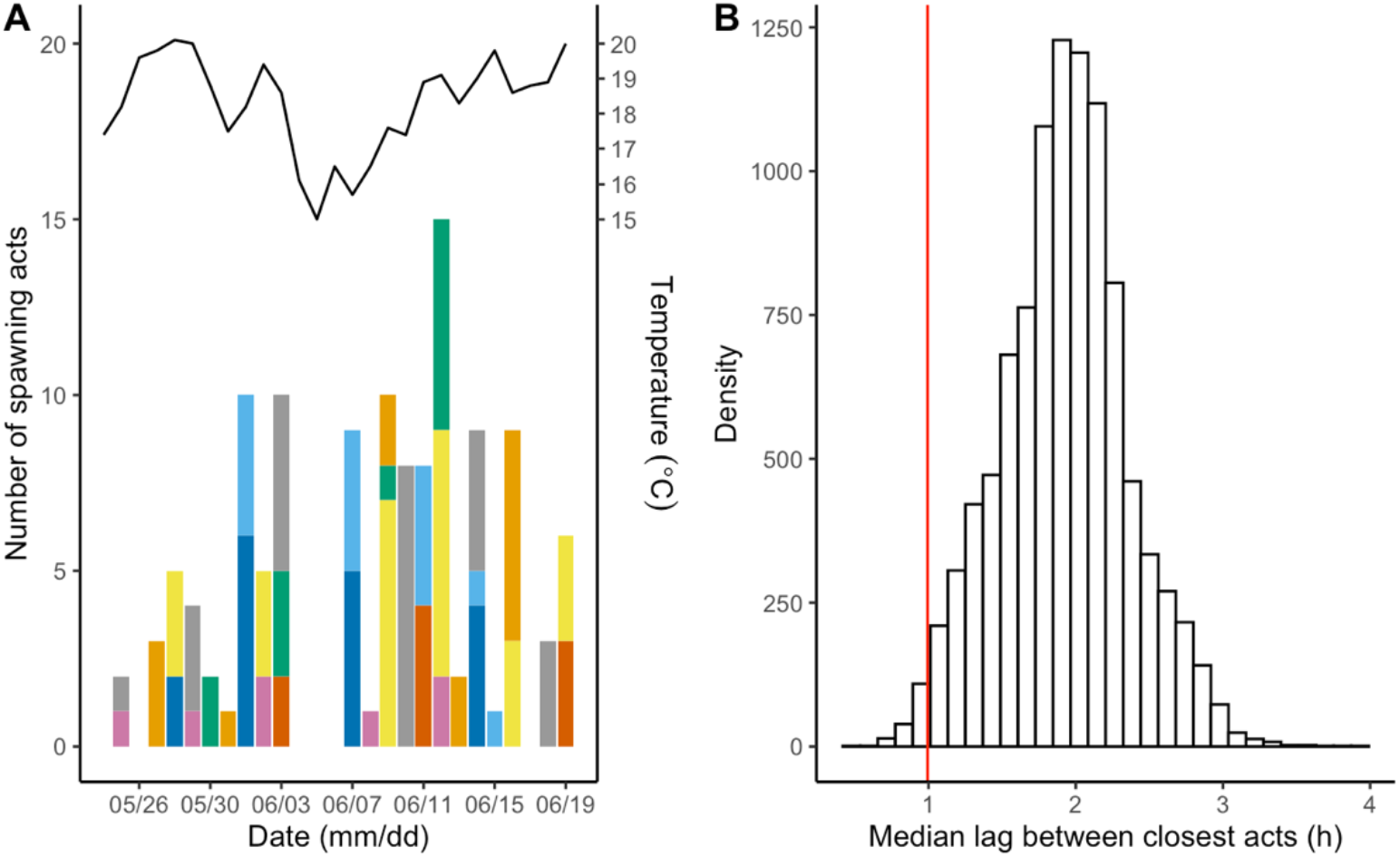
The temporal distribution of spawning acts by eight Allis shad females in the river Nivelle in spring 2017. **A.** Cumulated number of spawning acts for each night of the season. Each colour corresponds to an individual. The line above the bar plot represents the average water temperature measured each night between 10PM and 7AM. **B.** Synchrony of spawning acts performed by different females within a night (only the first act of the night, for each female). The red vertical line represents the median time lag between nearest spawning acts for observed data. The histogram represents the same thing for 10 000 permutation of the hour of the acts.

Average tail beat frequency (TBF), temperature and depth were computed for 312 840 minutes across all individuals’ spawning seasons (eight complete and one partial). Tail beat frequency ranged from 1.3 to 9.5 beats per second (mean=3.2), temperature ranged from 13.5 to 23.8°C (mean=18.3), and depth ranged from 0 to 400 cm (mean=186).

From equations (1) and (2), the estimated instantaneous rate of energy expenditure ranged from 0.09 to 0.89 kJ.min^−1^ (mean=0.17), and total energy expenditure by each of the eight individuals from initial capture to death ranged from 4 395 to 8 361 kJ (mean=6 277; Fig. 4.A). The corresponding mass loss was estimated to range from 127 to 241 g (mean=181), representing from 9% to 17% of initial mass (mean=12%). The shad died 44 to 182 hours (mean=98.62) after their last spawning act. They had lost between 33% and 53% of their mass (mean=42%), and their ovaries weighed 25.9 to 141.5 g (mean=79.7). No correlation was found between mass loss and ovary mass (Spearman S=89.3; rho=−0.06; p=0.888). The predicted and observed mass lost during the season were positively correlated, (Spearman S=8.55; rho=0.9; p=0.002; Fig. 4.B), but the observed mass loss was on average 3.7 times more than predicted (479 g difference on average).

**Figure 4.**
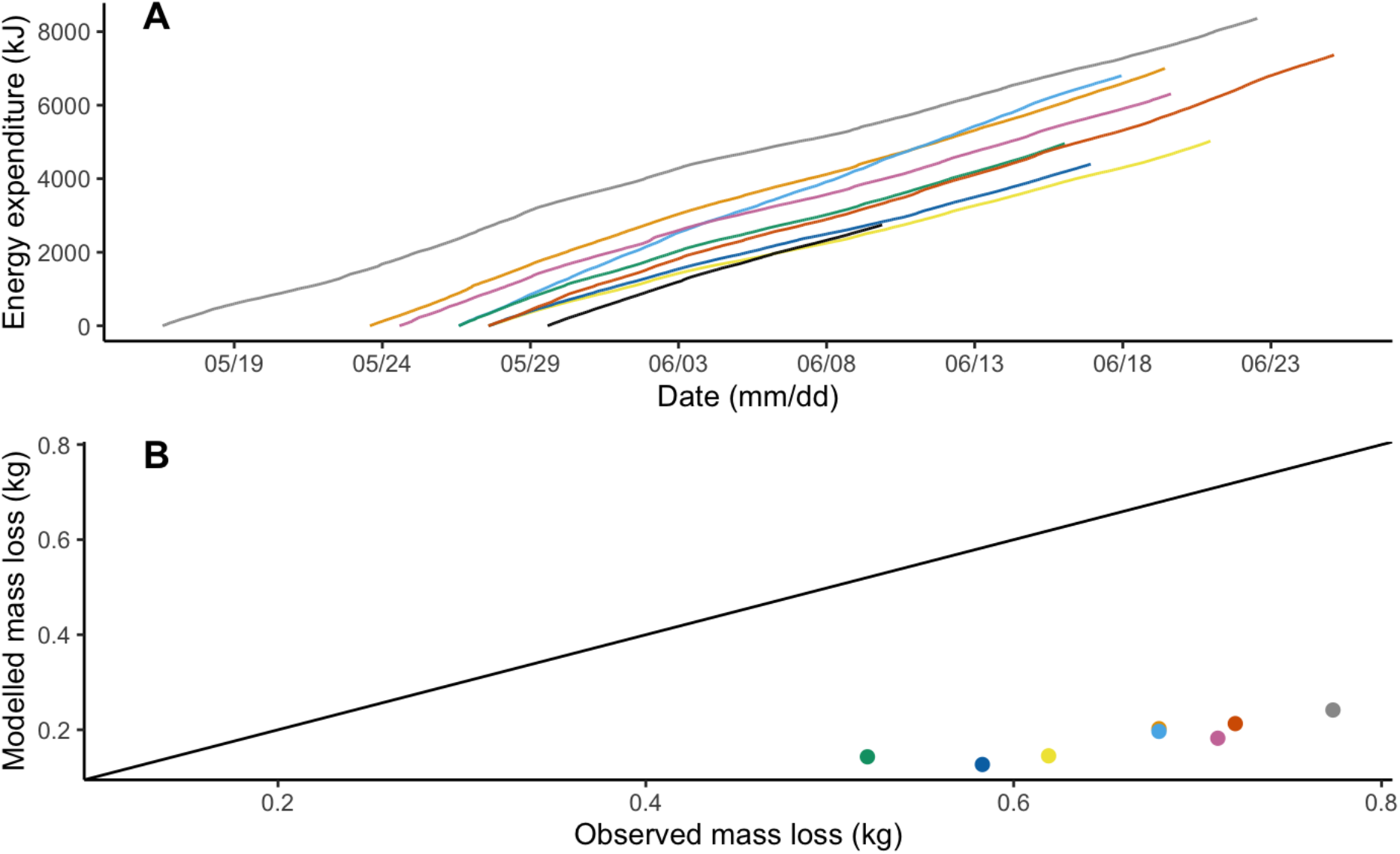
**A.** Energy expenditure over the spawning season for nine female Allis shad, estimated from temperature and tail beat frequency using equations (1) and (2). **B.** Observed mass loss and mass loss modelled with equations (1) and (2) over the spawning season. The straight line in B is the one on which points should lie if the modelled mass loss would fit the observations. Colours are as in Fig. 3, with black line in A for the individual whose tag stopped recording before the end of the experiment.

Aggregating TBF, temperature and pressure data over 8-hour periods (morning: 6AM-2PM, afternoon: 14PM-10PM, night: 10PM-6AM) produced 649 8-hour periods (Fig. 5). Shad stayed closer to the surface at night (mean ± standard error 160 ± 3 cm) than during the morning and afternoon (199 ± 3 cm) (LRT χ^2^=238.9; p<0.0001; R^2^_m_=0.14; R^2^_c_=0.69), and experienced warmer temperature in the afternoon (18.6 ± 0.1°C) than in the night (18.3 ± 0.1°C) than in the morning (17.9 ± 0.1°C) (LRT χ^2^=15.8; p<0.0001; R^2^_m_=0.02; R^2^_c_=0.05). TBF was higher at night (3.30 ± 0.03 beats/s) than in the afternoon (3.20 ± 0.04 beats/s) than in the morning (3.09 ± 0.03 beats/s) (LRT χ^2^=26.8; p<0.0001) and was also 0.21 beats/s higher during the first nine periods (three days) just after tagging than afterwards (LRT χ^2^=37.6; p<0.0001; R^2^_m_=0.06; R^2^_c_=0.36 for the model including both effects). The estimated energy expenditure was the highest at night (84 ± 1 kJ/8h), followed by afternoon (83 ± 1 kJ/8h) and morning (77 ± 1 kJ/8h) (LRT χ^2^=24.8; p<0.0001; R^2^_m_=0.03; R^2^_c_=0.34). The mass of eggs remaining at death was not correlated to the energy expenditure cumulated across mornings (Spearman S=66; rho=0.21, p=0.619), afternoons (Spearman S=64; rho=24; p=0.582), nights (Spearman S=76; rho=0.09; p=0.84), or the whole season (Spearman S=64; rho=0.24; p=0.582).

**Figure 5.**
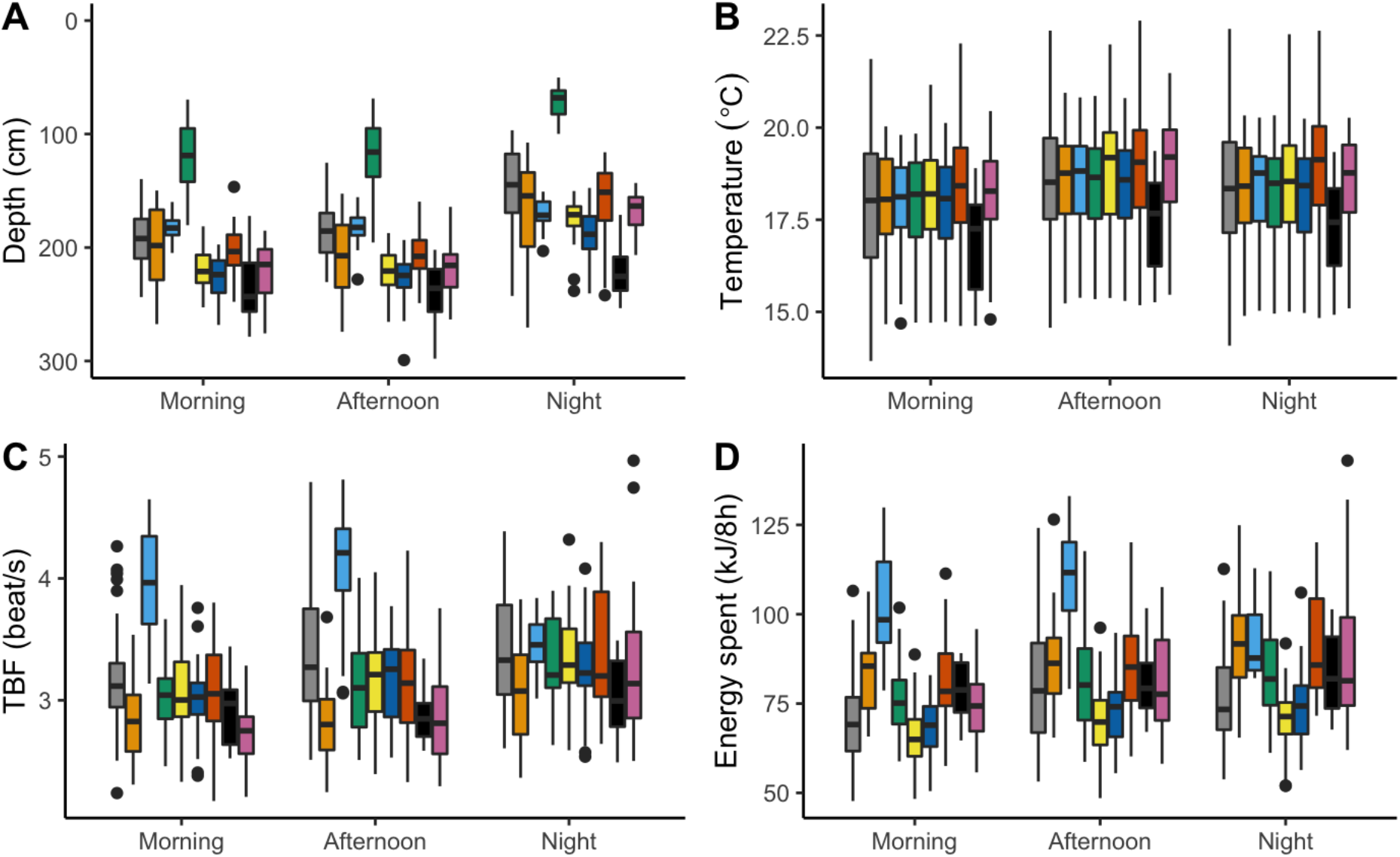
Depth (**A**), temperature (**B**), tail beat frequency (**C**) and estimated energy spent (**D**) by nine female allis shad during the morning (6AM:2PM), afternoon (2PM:10PM) and night (10PM:6AM) of their spawning season. Box plots show the distribution of the variables averaged (A, B, C) or cumulated (D) across 8-hour periods for each individual (colours as in Fig. 3, black for the individual whose tag stopped recording before the end of the experiment). In C, energy spent was computed from temperature and TBF using equations (1) and (2).

As expected, the *θ* angle between the x-axis of the accelerometer and the vertical plane was positively correlated to the individual’s Fulton coefficient of condition (Pearson’s r=0.63; p=0.006), although the effect seemed to be due to within-individual difference in condition and *θ* at the beginning and at the end of the season rather than inter-individual variability in condition and *θ* (Fig. 6.A). The *θ* angle globally decreased over time for all individuals and was negatively related to depth, resulting in occasional increases in *θ* during 8-hour periods when shad stayed closer to the surface (full model R^2^_m_=0.28; R^2^c=0.79; Fig. 6.B). The effect of time corresponded to a slope of −0.63 degree per day (LRT χ^2^=518.4; p<0.001) and the effect of depth corresponded to a slope of −0.017 degree per cm (LRT χ^2^=9.6; p=0.002). Contrary to our expectation, the difference of the *θ* angle between initial capture and death was not related to change in body condition between initial capture and final recapture (Spearman S=76, rho=0.09, p=0.84). The best mixed model to explain the shift in *θ* during a 8-hour period was the one with the moment (morning, afternoon or night) of the period as the independent factor (LRT χ^2^=56.7; p<0.0001; R^2^_m_=0.08; R^2^_c_=0.08), followed by the null model including individual random effect only (ΔAICc=51), the model including estimated energy expenditure as the independent variable (ΔAICc=56), and the model including both temperature and TBF as independent variables (ΔAICc=58). According to the best model, the shift in *θ* was negative during mornings (mean ± standard deviation −1.25° ± 3.08), slightly negative during afternoons (−0.23° ± 2.36), and positive during the nights (0.89° ± 3.15).

**Figure 6.**
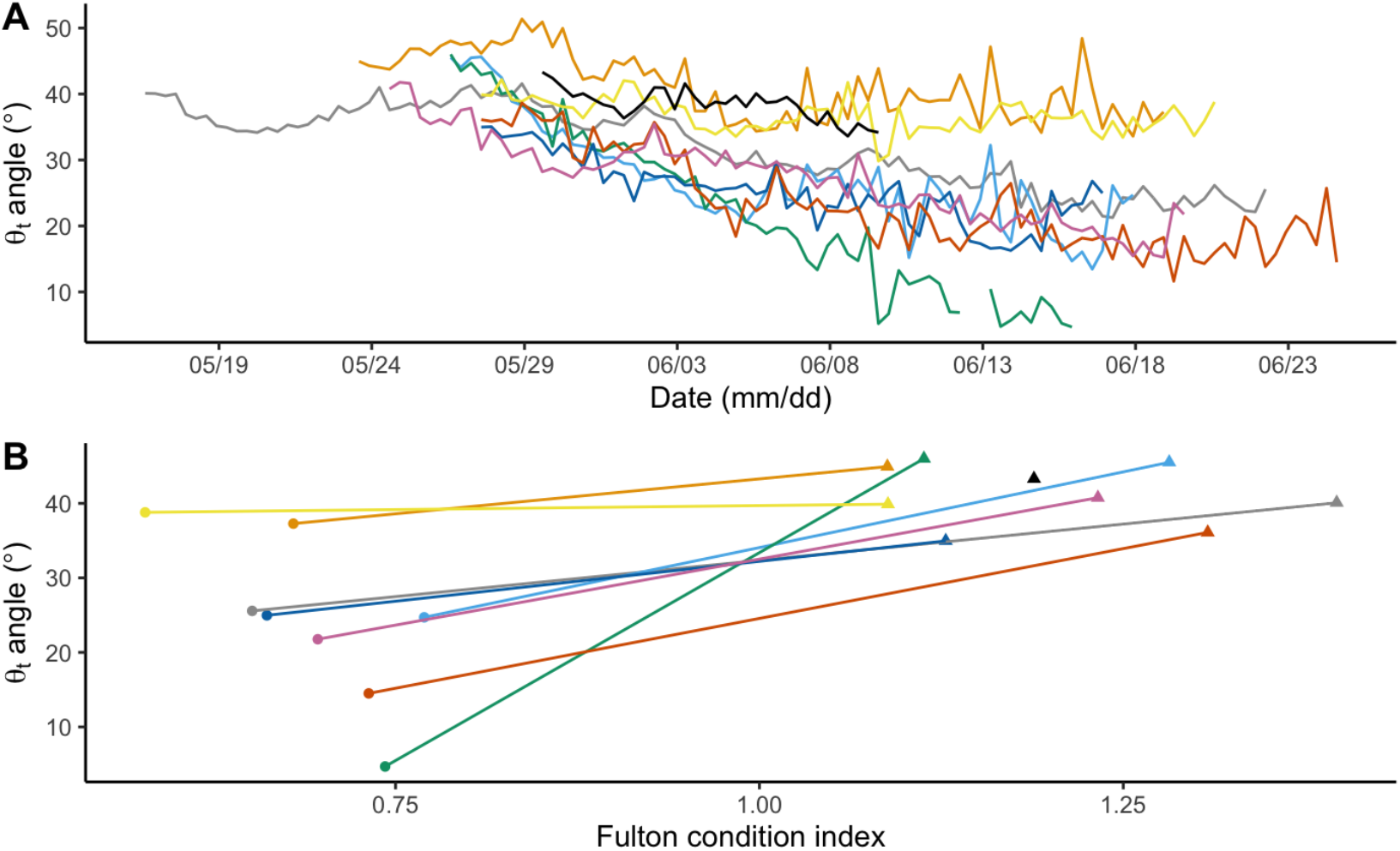
The *θ_t_* angle between the x-axis of the accelerometer and the vertical plane as an indicator of fish body roundness. **A**. Temporal evolution of *θ* computed each day along the spawning season for nine Allis shad. **B.** Relationship between Fulton coefficient of condition and *θ_t_* angle at initial release (triangles) and at recapture after death (points). Each individual is represented with the same colour as in Fig. 3, and black is for the individual whose tag stopped recording before the end of the experiment.

## Discussion

In this study, we used acceleration data to quantify the dynamics of spawning acts, energy expenditure and slimming of female Allis shad throughout their spawning season. The results indicate that the spawning schedule of female shad, although constrained by the serial maturation of oocyte batches, was also influenced by temperature and social factors: females had a higher probability to spawn during warmer nights, spawned repeatedly during most of their active nights and females that spawned on the same nights synchronized their spawning acts with each other. Tail beat frequency and water temperature recorded by the loggers showed that energy expenditure was slightly higher during nighttime than during daytime, and may have been so high that shad died from energy exhaustion before having spawned all their eggs. Finally, the novel use of gravitational acceleration to monitor fish slimming, although perfectible, seems a promising method to monitor changes in animal condition in the field.

### Dynamics of spawning acts and energy expenditure

The number of spawning acts performed by female Allis shad ranged from 7 to 26, with an average of 15.75. So far, the average individual number of spawning acts was indirectly estimated to be either five (Acolas et al., 2006) or 12 (Fatin and Dartiguelongue 1996) by counting the number of acts heard upstream a dam where all passing individuals were censused. Such method provides no estimate of interindividual variability. In a first attempt to work at the individual level, Acolas et al. (2004) marked three females and three males with acoustic tags, and based on the number of detections of the tags near the surface (assessed by the reception power by hydrophones immersed at different depth), estimated that zero, one and two acts were performed by the females and three, 38 and 60 acts by the males. Here, despite a small sample, we estimated a more representative distribution of the number of acts per individual, as their detection on accelerograms is certainly more reliable than the method used by Acolas et al. (2004). This distribution, as well as the timing of spawning acts, is crucial to build a model estimating the number of shad in a river from the acoustic survey of spawning acts, as it is routinely done in European rivers (e.g. Chanseau et al. 2004). Such a model, simulating shad spawning behaviour in an Approximate Bayesian Computation framework (ABC; Csilléry et al. 2010) is available in French here: https://ctentelier.shinyapps.io/alose_abc/. The number of acts per female could be correlated to none of the few biometric variables collected, but the very small sample implied a weak statistical power. Larger females, which have more eggs and more energy, did not perform more spawning acts (a nonsignificant tendency towards negative correlation was observed).

Beside oocyte maturation and temperature, the temporal distribution of spawning acts was aggregated within and among females. While each female was active for most of the population’s spawning season, the individual spawning season was punctuated by three to six nights of activity generally corresponding to volleys of two to eight acts performed in a few tens of minutes, and separated by on average 3.5 nights of inactivity. The schedule of spawning acts must be constrained by the fragmented maturation of oocytes (Cassou-Leins and Cassou-Leins 1981; Olney et al. 2001), but also depended on temperature. Interestingly, while data collected at the population level indicate that spawning activity increases with temperature (Paumier et al. 2019), our data collected at the individual level showed that temperature increased the probability that a female performed some acts during the night (the zero-inflation component of the zero-inflated regression), but not the number of acts it performed (the count component of the zero-inflated regression). In fact, two results suggest that the dynamics of spawning acts within a night might be influenced by social factors. First, the temporal proximity of acts performed by a given female in a given night in this study, and the high proportion of acts performed without oocyte expulsion reported by Langkau et al. (2016) are reminiscent of female trout’s ‘false orgasm’ (Petersson 2001) or female lamprey’s ‘sham mating’ (Yamazaki and Koizumi 2017). For both of these species, it has been suggested that repeated spawning simulations enable the female to exert mate choice, by both exhausting the sperm stock of an unwanted courtier and signalling its mating activity to peripheral males. Second, spawning acts of different females in a given night were more synchronous than expected from the hour of spawning acts across all nights of the season. This suggests that a female which is ready to spawn (mature batch of oocytes) in a propitious night (warm temperature) may trigger its spawning acts when another female does so. Such fine scale synchrony in the mating activity of different females may again affect sexual selection, reducing the environmental potential for polygyny by making it difficult for the same male to monopolize several females at the same time (Emlen and Oring 1977).

The end point in the spawning schedule of semelparous organisms is death, which occurred in female shad two to seven days after their last spawning act, and before they laid all their eggs, which suggests that energetic reserves were exhausted before egg stock. Combining acceleration and temperature data collected in the field with equations parametrized in laboratory experiments, we estimated that the energetic expenditure of spawning shad during three weeks of spawning ranged from 4 395 to 8 361 kJ (mean=6 277). This is surprisingly of the same order as female American shad, which entered the Connecticut river with a stock of 12 000 kJ, of which approximately 5 000 kJ were consumed along their 228-km and seven-week long upstream migration in waters warming from 10 to 22°C (Leonard and McCormick 1999; but see next section for possible biases in our estimation). Extreme pre-spawning energy expenditure caused by long migration, obstacle crossing or warm water can reduce the reproductive lifespan of semelparous females and hinder their ability to spawn their full egg stock (Hruska et al. 2011). While the spawning ground used by shad in our study is only 13 km from the river mouth with only one obstacle to pass (Uxondoa, where we captured them), the median daily temperature recorded in the Nivelle for May and June 2017 was 18.4°C, the second warmest spring since the beginning of record in 1984, the maximum being 18.5°C in 1989, the minimum 14.6°C in 2013, and the median 16.9°C. According to our analysis of data from Leonard et al. (1999; our equation 1), an increase in temperature from 16.9 to 18.4°C would raise the energy expenditure by 8.3%, giving both a possible explanation for the inability of shad to spawn all their eggs before they die and a hint of the impact of global warming on the breeding performance of such species.

As expected from visual observations on spawning grounds, shads were more active at night than during the day, and stayed deeper during daytime when water was warmer. Although nocturnal spawning is usually interpreted as a strategy to decrease predation risk on spawners or eggs (Robertson 1991), resting in deeper, hence fresher, water during the warm hours of the day may also save energy for spawning at night. However, contrary to our predictions, the individuals that spent less energy during mornings or afternoons and more during the night did not have fewer residual eggs at death. Although differences between mornings, afternoons and nights were tenuous and our small sample size precludes conclusive inferences, the continuous measurement of spawning activity and energy expenditure at the individual level opens the door to the quantification and decomposition of interindividual variation in the schedule of reproductive behaviour and energy management. Moreover, coupling telemetric data on behaviour with additional fitness indicators such as offspring production (egg sampling and genetic parentage analysis) could inform on the strength of natural and sexual selection acting on these phenological strategies in the field (Tentelier et al. 2016).

### Methodological considerations

According to calculated energy expenditure, the energetic model predicted that shad should have lost on average 12% of their initial mass, which was much less than the observed 42% loss. Of course, the energetic model did not account for egg expulsion, which must represent a large proportion of mass loss, given that ovaries can represent up to 15% of somatic mass at the onset of the spawning season (Cassou-Leins and Cassou-Leins 1981; Taverny 1991). Moreover, the equations used to convert TBF and temperature to energy expenditure and mass loss were not parametrized with data obtained on *A. alosa* but on *A. sapidissima* (Castro-Santos & Letcher, 2010; Leonard et al., 1999; Leonard & McCormick, 1999) which is the species most closely related to *A. alosa* for which such reliable information exist. Although American shad and Allis shad have the same morphology and ecology, some elements suggest that results on American shad could not exactly apply to our study on Allis shad, and that our estimates of energy expenditure should be taken cautiously. First, although the range of temperatures used by Leonard et al. (1999) were similar to the temperatures encountered by shad in our study, TBF of Allis shad in the field exceeded the fastest swim tested by Leonard et al. (1999). In particular, the brief bouts of very high TBF which correspond to spawning acts may represent anaerobic efforts, which are sustained by white muscle rather than red muscle and are more costly than the aerobic effort observed by Leonard et al. (1999). Given the difference in mechano-chemical efficiency (ratio of mechanical power output to chemical energy consumed) between red and white muscles, a different equation should be calibrated for each type of effort (Gleiss et al. 2011). The volleys of spawning acts performed by females in a few tens of minutes could even lead to sexual selection of traits affecting the recovery after sprint (Kieffer 2000). Second, the relationship linking swim speed to oxygen demand (MO2) varies between species, even closely related, and American shad seems to have a particularly high metabolism compared to other clupeids (Leonard et al. 1999). This would have led to an overestimation of energy costs for *A. alosa*. Third, the relative contribution of lipids and proteins as metabolic fuel may differ between the iteroparous American shad and the semelparous Allis shad. Indeed, as migratory iteroparous fishes have to conserve proteins, especially in the red muscle, for their downstream migration, semelparous species can exhaust their protein stock to complete spawning (Schultz 1999). Also, Leonard & McCormick (1999) measured the relative expenditure of lipids and proteins in *migrating* American shad, while we monitored *spawning* Allis shad. It has been shown that most anadromous fishes catabolise proteins only when lipid reserves have been exhausted, in a way that lipids mainly fuel migration while proteins fuel spawning (Idler and Bitners 1958; Beamish et al. 1979; Jonsson et al. 1997; Hendry and Berg 1999). Given that proteins have a lower energy density than lipids, the surprising high energy consumption and the low mass loss predicted by the energetic model may be explained by spawning Allis shad relying more on proteins than on lipids. All these caveats could be avoided by laboratory calibration of the equations relating energy expenditure to tail beat frequency and temperature on the same species, at the same stage of energy reserves, bearing the same tags and performing the same behaviours as in the field.

While the presence of the radio tag and the accelerometer may have imposed an energetic cost on shad due to additional mass (less than 2% of initial fish mass) and drag, several indicators suggest that our tagging method did not impact shad behaviour as heavily than the commonly gastric implants which result in high mortality rate or long post-tagging downstream movements (Steinbach et al. 1986; Verdeyroux et al. 2015; Tétard et al. 2016). Such downstream movements were only observed in 2018, after exceptional floods which have probably also make untagged shad drift downstream, especially after the spawning season. Although the dead individuals we retrieved in 2017 showed clear depigmentation, neither severe abrasion nor fungal proliferation was observed, even after one month in water at 18° C. Moreover, the eight females for which complete accelerograms were retrieved all spawned, three of them spawned the night directly following tagging, and the median of 14 spawning acts was above the five to twelve acts per individuals estimated by Acolas *et al.* (2006) and Fatin & Dartiguelongue (1996). However, shad may have suffered post-tagging stress, as indicated by the higher tail beat frequency (TBF) during the three days following tagging that during the remainder of the spawning season, similar to the increased activity of Atlantic salmon during the three days after tagging (Føre et al. 2020). On the other hand, a higher TBF just after tagging may be due to the fish finishing their upstream migration, before settling near spawning grounds (unpublished radio tracking data). This moderate negative impact suggests that external tagging under the dorsal fin, provided fish are continually kept immerged and rapidly handled, is a suitable tagging technique for Allis shad (Jepsen et al. 2015; Breine et al. 2017). Although the swimming behaviour did not seem to be strongly impaired, the additional weight or drag force caused by the tag may have increased the energy required to perform this behaviour (Jepsen et al. 2015). In this case, the energy expenditure of tagged fish computed from equations derived from swim tunnel experiments performed on untagged fish would be underestimated. To develop the telemetric measurement of realistic energy expenditure, attention should be paid to both designing tags that do not impose too much additional energetic costs on individuals and estimating these additional costs.

On top of the estimation of energetic costs from dynamic acceleration, the continuous estimation of body roundness from static acceleration is a promising yet perfectible application. Gravitational acceleration is commonly used to infer the posture of the animal, assuming that the position of the accelerometer on the animal is constant (Brown et al. 2013). Here, assuming that shad stayed upright for most of the time, gravitational acceleration was used as an indicator of change in body shape. In particular, we hypothesized that the angle *θ* between the *x* axis of the accelerometer and the vertical plane may indicate body roundness. For all individuals, *θ* was higher at the beginning of the season than at the end, when the fish had slimmed, and globally decreased along the spawning season. The accelerometer was attached under the dorsal fin, where the cross-section is the broadest, which probably maximized the sensitivity of acceleration data to fish slimming. Furthermore, shad mainly store energy for migration and spawning as subdermal fat and interstitial fat in the white muscle (Leonard and McCormick 1999), whereas the limited consumption of visceral and liver fat may not have affected *θ*. However, interindividual variability in *θ* was not correlated to interindividual variability in body condition, and the shift in *θ* was not correlated to individual mass loss during the spawning season. This could be due to interindividual differences in the exact position of the accelerometers on the fish, or in the relationship between body condition and roundness. Indeed, mass loss due to gamete expulsion was probably undetectable by the accelerometer attached to the back of the fish. Given that ovaries can represent up to 15% of a shad mass at arrival on spawning grounds (Cassou-Leins and Cassou-Leins 1981; Taverny 1991), a significant loss of mass was probably not reflected in the rotation of the accelerometer across the spawning season. In fact, shift of *θ* across eight hour periods increased with neither TBF, temperature, nor estimated energy expenditure, contrary to what was expected from the relationship linking TBF and temperature to energy consumption in American shad (Leonard et al. 1999). This lack of resolution may be due to the way tags were attached. In particular, while the attachment wires were tightened during the tagging procedure, fish slimming loosened them so the accelerometer must have jiggled more and more as fish slimmed, thereby increasing the noise in acceleration data. Given all these results and limits, using the angle of the accelerometer to quantitatively track change in mass or body roundness will certainly require further tuning, such as laboratory experiment with repeated weighing and image analysis of the fish’s cross-section, and drastic standardization of attachment procedure, but we consider it a promising method to monitor individual condition in the field. On top of slimming, *θ* was linked to depth, and increased during some periods when the fish stayed closer to the surface, in particular during nights. This was certainly due to the inflation of the swim bladder, which represents around 8% of the volume of the fish, and inflates as the fish stays closer to the surface (Alexander 1966). Although here, swim bladder inflation introduced additional variation in our monitoring of fish slimming, it could also be the process of interest in some field studies, for example to untangle the contribution of swim bladder inflation and active swimming in the vertical migration of pelagic fishes (e.g. Pelster 2015). This illustrates the potentially broad application of our acceleration-based approached, which, once refined, could be used to detect the many ecologically or behaviourally relevant changes in animal shape beyond slimming due to energy consumption, such as parturition, massive food intake in large predators (Cuyper et al. 2019), inflammatory swelling (Duncan et al. 2016), or inflation as a courtship or defence behaviour (Wainwright and Turingan 1997). Combined with dynamic acceleration, such data could be used to test whether these changes in shape are associated to global activity or to the expression of specific behaviours.

## Data accessibility

Data are available online: https://doi.org/10.15454/NTFYCC.

## Supplementary material

Script and codes are available online: https://doi.org/10.15454/NTFYCC.

## Acknowledgements

We thank B. Liquet, F. Luthon, B. Larroque and E. Bouix for their help in the initial exploration of data, and J. Leonard and T. Castro-Santos for sharing their data on American shad. This study was financed by the Adour-Garonne Water Agency, and the French region Nouvelle Aquitaine. Version 7 of this preprint has been peer-reviewed and recommended by Peer Community In Ecology (https://doi.org/10.24072/pci.ecology.100060)

